# Depletion of erythropoietic miR-486-5p and miR-451a improves detectability of rare microRNAs in peripheral blood-derived small RNA sequencing libraries

**DOI:** 10.1101/789891

**Authors:** Simonas Juzenas, Carl Mårten Lindqvist, Go Ito, Yewgenia Dolshanskaya, Jonas Halfvarson, Andre Franke, Georg Hemmrich-Stanisak

**Affiliations:** Institute of Clinical Molecular Biology, Christian-Albrechts-University of Kiel, DE 24105 Kiel, Germany; School of Medical Sciences, Faculty of Medicine and Health, Örebro University, SE 70182 Örebro, Sweden

## Abstract

Erythroid-specific miR-451a and miR-486-5p are two of the most dominant microRNAs (miRNAs) in human peripheral blood. In small RNA sequencing libraries, their overabundance reduces diversity as well as complexity and consequently causes negative effects such as missing detectability and inaccurate quantification of low abundant miRNAs. Here we present a simple, cost-effective and easy to implement hybridization-based method to deplete these two erythropoietic miRNAs from blood-derived RNA samples. By utilization of blocking oligonucleotides, this method provides a highly efficient and specific depletion of miR-486-5p and miR-451a, which leads to a considerable increase of measured expression as well as detectability of low abundant miRNA species. The blocking oligos are compatible with common 5’ ligation-dependent small RNA library preparation protocols, including commercially available kits, such as Illumina TruSeq and Perkin Elmer NEXTflex. Furthermore, the here described method and oligo design principle can be easily adapted to target many other miRNA molecules, depending on context and research question.

## Introduction

Small RNA sequencing (smRNA-Seq) is a widely used application, enabling discovery and quantification of small RNAs, including microRNAs (miRNAs), which regulate gene expression and are emerging as important disease biomarkers (1, 2). Many of the current miRNA-based biomarker studies use plasma, serum, extracellular vesicles (EVs) or peripheral blood as a source material. The biggest advantage of peripheral blood over the other blood-derived biological materials comes from very low technical variability during sample handling and processing, which makes this material very attractive for the use in a clinical setting (3). Moreover, peripheral blood contains complete convolved information about miRNA expression of all cellular and non-cellular blood compounds. On the other hand, some of these compounds in whole blood are causing more problems than benefits. For example, in addition to globin mRNAs, red blood cells also contain highly abundant miRNAs including miR-486-5p and miR-451a (4, 5). These two conserved, non-canonical miRNA molecules represent the most dominant miRNAs in peripheral blood, which are specifically upregulated in the erythroid lineage (6). Loss of *mir-486* and *mir-451a* genes in mice leads to erythroid defects, showing their importance in erythrocyte development (6), and explaining their high abundance in erythroid cells.

In RNA-Seq workflows, significant overabundance of transcripts drastically reduces the complexity of PCR-amplified sequencing libraries and exhausts sequencing space, which leads to unwanted effects, such as inaccurate detectability and quantification of low abundant transcripts. Especially in the biomarker field, low level of detectability is a very critical issue, since many disease-derived miRNAs circulate in blood at low amounts (7).

Concerns about depletion of unwanted miRNAs in small RNA libraries are not new, and at least two approaches have been described previously (8, 9). The hybridization-based approach relies on stem-loop shaped oligonucleotides with a 12-base 5’ overhang which represents the reverse complement to the first 12 bases of the 5’ end of the target miRNA’s canonical sequence (8). The other method is a CRISPR/Cas9-based approach, where Cas9 is complexed with single guide RNAs that target undesirable miRNA sequences for cleavage *in vitro* (9). Both methods have been shown to effectively reduce target miRNA sequences in small RNA libraries; however, both of them have their own limitations. Whereas the hybridization-based approach has a high chance of unspecific hybridization with non-target miRNAs due to the short complementarity (12 bases) of the stem-loop oligonucleotide, the CRISPR/Cas9-based method includes additional steps and reagents which increase hands-on-time and costs of the procedure.

Besides unwanted miRNA depletion, there are more methods developed to remove other single stranded nucleic acids from small RNA-sequencing libraries, i.e. tRNA-derived small RNA depletion using biotinylated DNA probes together with streptavidin coated magnetic beads or DNA probes with RNase H and DNase 1 (10); rRNA depletion using DNA oligonucleotides having 3’ C3 (11) and 3’ biotin (12) terminal modifications; adapter-dimer depletion using LNA-modified oligos (13).

Here, by employing DNA oligonucleotides, we describe a simple, cost-effective and easy to implement hybridization-based protocol for miR-486-5p and miR-451a depletion from small RNA sequencing libraries. The method involves linear oligonucleotides covering selected target miRNA sequence variants to block 5’ adapter ligation. In addition, the described method is compatible with a wide range of commercially available 5’ ligation-dependent small RNA library preparation protocols.

## Methods

### Blood miRNA catalog data analysis

Compositional analysis of miRNA species in blood compounds was performed using processed miRNA count data from our previously published study (14). The dataset contains miRNA expression counts of seven types of blood cells, EVs, serum and whole blood, which were generated by using TruSeq (Illumina) small RNA library preparation protocol. The miRNA read counts for each sample in the dataset were down-sampled to a constant number of reads using random subsampling rrarefy() function implemented in the R package vegan (15). For visualization, relative abundance for each miRNA (i) in each sample (j) was estimated using the following formula:

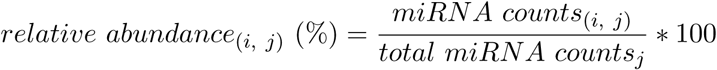

Shannon diversity index was calculated for each sample based on down-sampled miRNA counts by employing the diversity() function from the R package vegan (15). A lower-tailed t-test was performed to evaluate the differences of the index values between whole blood and every other blood compound using t.test() function from the base R package.

### Blocking oligo design

Erythropoietic miR-486-5p and miR-451a blockers are linear single-strand DNA oligonucleotides with 3’ C3 spacer (propyl group) modification (IDT). The oligos are designed to prevent 5’ adapter ligation via blocking the access of T4 RNA ligase 1 to the 5’ end of target miRNAs during smRNA-seq library preparation. Sequences of two blocking oligos were composed based on our previously published whole blood smRNA-seq data (14). Briefly, all sequences that were mapped to precursors of miR-486-5p and miR-451a were pooled in order to obtain all possible unique sequence variants for each miRNA. The obtained unique sequences for each target miRNA were then flattened into a single consensus sequence in order to retrieve the most frequent nucleotide found at each position in a sequence alignment. The consensus sequences were then used to generate reverse complement oligonucleotides to bind target miRNAs by Watson–Crick base pairing. The C3 spacer modification was attached to the 3’ ends of oligonucleotides to avoid self-ligation. The resulting oligonucleotides are provided in **Figure 2A**.

**Figure 1:**
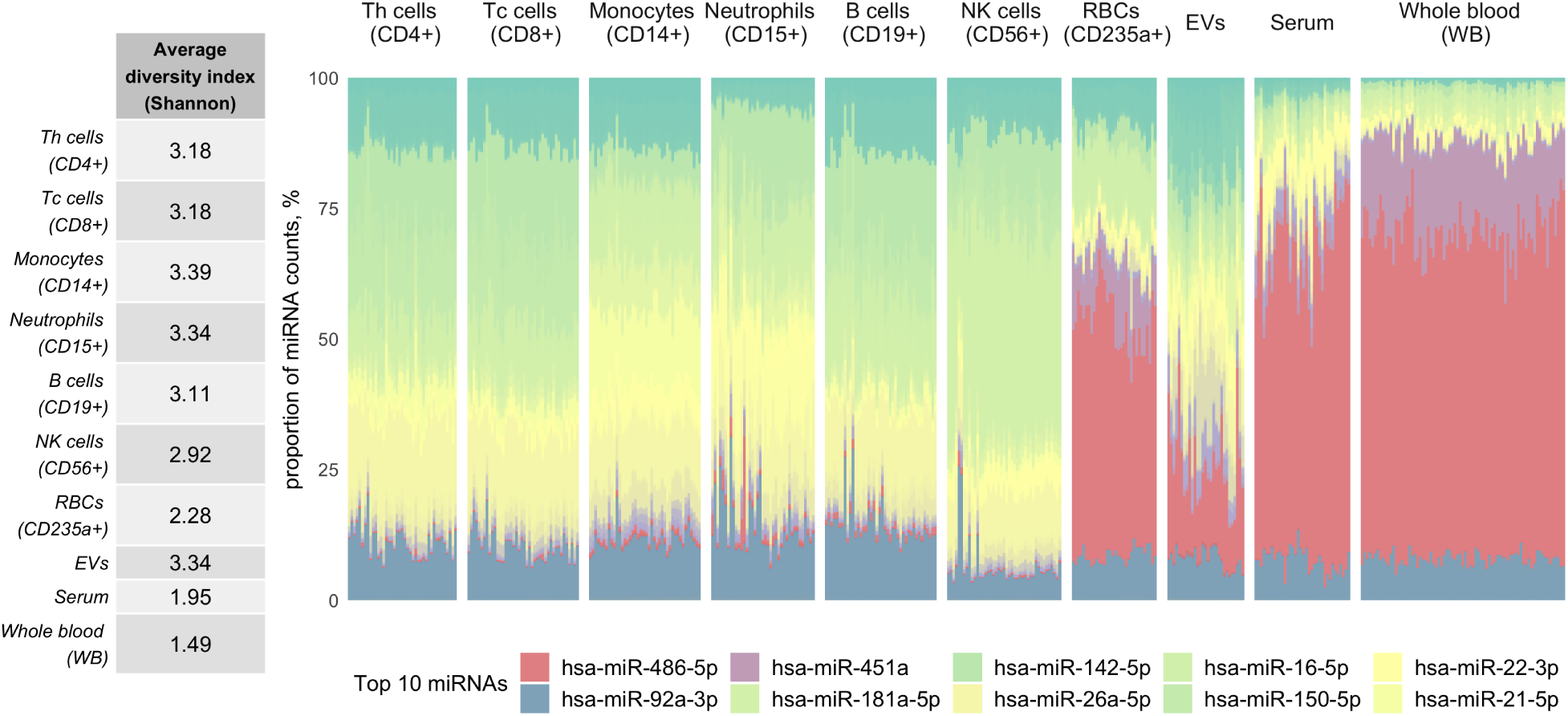
Erythropoietic miR-486-5p and miR-451a domination reduces diversity of whole blood-derived small RNA libraries. The left side panel represents averaged Shannon diversity index of every blood compound. Shannon index value for each sample was calculated on down-sampled miRNA count data, and then the values were compared between whole blood and every other compound using pairwise t-test. The significantly lowest miRNA diversity is observed in whole blood (PAXgene) samples. In the right side panel, a stacked bar chart presents relative abundance values of miRNAs in different blood compounds, where every bar represents a sample and every color represents a fraction of single miRNA within a given sample. The graph reveals high dominance of erythropoietic miR-486-5p and miR-451a in whole blood, red blood cell (RBC), serum and extracellular vesicle (EVs) samples. The ten most abundant miRNAs in all of the blood compounds are shown in the legend.

**Figure 2:**
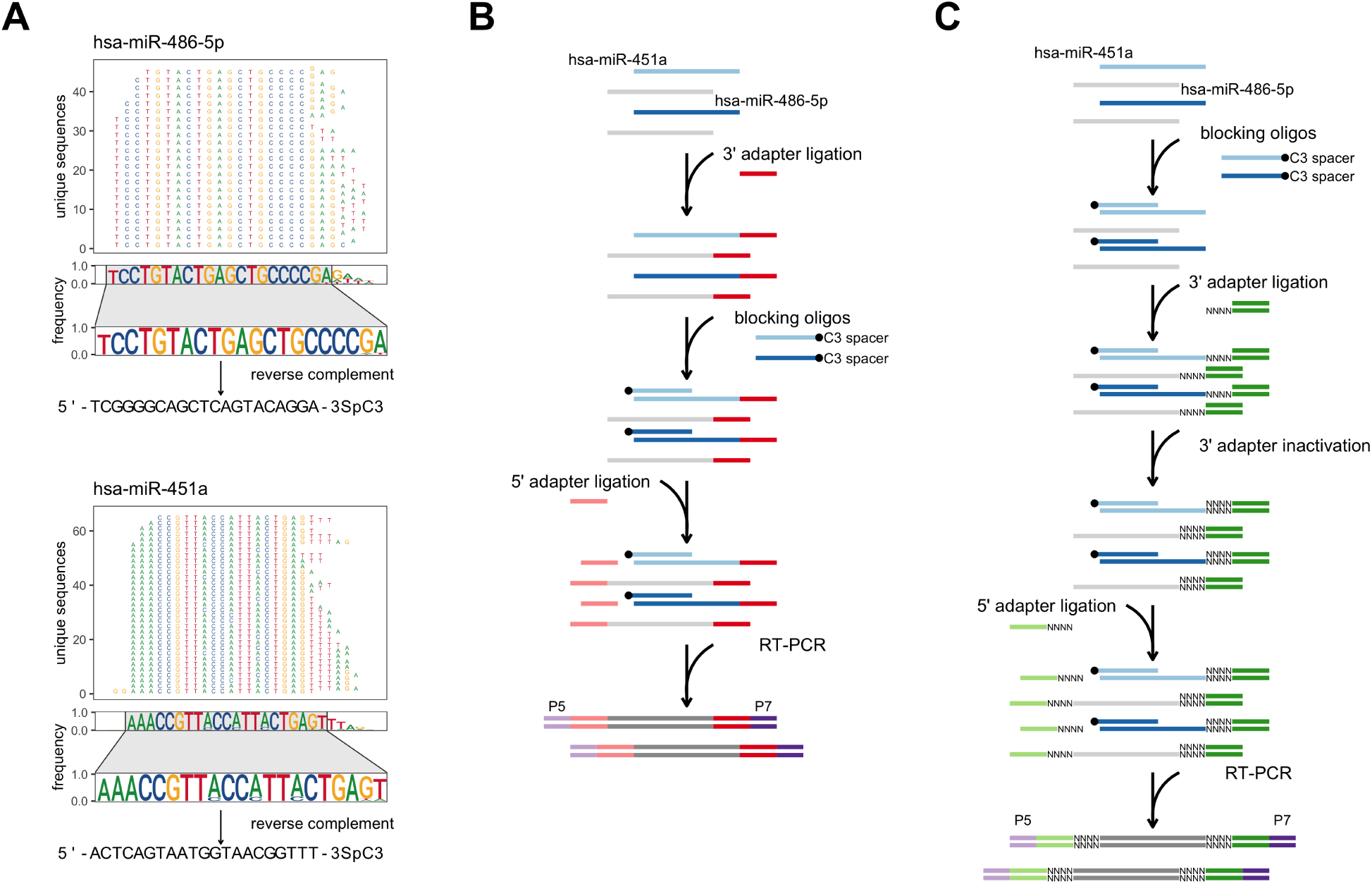
Blocking oligo design and application workflows using modified TruSeq and NEXTflex small RNA library preparation protocols. (**A**) Design principle of miR-486-5p and miR-451a blocking oligonucleotides. Briefly, unique pooled-sample sequences which mapped to precursors of miR-486-5p and miR-451a were used to retrieve the most frequent nucleotide found at each position in a sequence alignment. The most stable consensus sequences were used to generate reverse complement DNA oligonucleotides of targeted miRNAs. The C3 spacer (propyl group) modification was added to the 3’ ends of the synthetic oligonucleotides to avoid self-ligation. Whole blood smRNA-seq data used for oligo design was obtained from GSE100467; (**B**) A schematic representation of modified Illumina’s TruSeq small RNA library preparation protocol. The modified protocol involves an additional step, where synthetic blocking oligonucleotides are introduced right before the 5’ adapter ligation reaction. In this step, the blocking oligonucleotides are annealed to target miRNAs, which results in double-stranded RNA:DNA hybrid formation. These blunt-ended or slight 3’ DNA overhang-having double-stranded hybrids are not suitable substrates for T4 RNA ligase-mediated addition of adapter oligonucleotide to the 5’ end of RNA strand in the hybrid. As a consequence, blocked RNA:DNA hybrids without 5’ adapter sequences cannot be amplified and therefore are depleted from final small RNA library; (**C**) A schematic workflow of modified Perkin Elmer’s NEXTflex small RNA library preparation protocol. In comparison to TruSeq, the standard NEXTflex protocol includes an extra step called 3’ adapter inactivation, where end-filling is performed to fill the gaps of random nucleotides bearing 5’ overhang portions of 3’ adapter duplexes. Because of this step, in order to avoid denaturation of the 3’ adapter duplexes, blocking oligos for miR-486-5p and miR-451a were introduced directly to total RNA sample.

### RNA extraction

Whole blood samples from 10 healthy volunteers (n = 5 females and n = 5 males) aged from 25 to 37 were collected into PAX gene RNA blood tubes (Qiagen). Total RNA samples were isolated using QIAcube automation with the PAXgene Blood miRNA Kit (Qiagen) in accordance to manufacturer’s instructions.

### Small RNA-seq and erythropoietic miRNA blocking

Small RNA libraries were prepared using standard and modified TruSeq Small RNA Library Prep Kit v02 (Illumina) and gel-free NEXTFLEX Small RNA Seq Kit v3 (Perkin Elmer) protocols. The modified versions of the protocols include an additional step where blocking oligos for miR-486-5p and miR-451a are included. In case of the TruSeq protocol, 1 *µ*l of 20 *µ*M of the blocking oligo mix was introduced and annealed immediately after the 3’ adapter ligation (ramp from 65 to 45 ° C – 0.1 per sec), whereas for the NEXTflex protocol, 1 *µ*l of 10 *µ*M of the blocking oligo mix was introduced and annealed directly to the total RNA sample prior to library preparation (detailed protocols are provided in **Supplementary Methods**). In order to achieve the best performance of the TruSeq and NEXTflex library preparation methods, starting RNA amounts were selected to be in a range of manufacturer’s provided recommendations. Concisely, for each TruSeq (standard and modified) library preparation, 1 *µ*g of total RNA was used as starting material, whereas for each NEXTflex (standard and modified) library preparation, 100 ng of total RNA was used as input. Subsequently, for each sample standard and modified libraries were generated and randomized in a supervised fashion (blocked and unblocked paired samples on the same lane to minimize batch-effect) and pooled with 10 samples per lane. Sequencing was performed on an Illumina HiSeq 4000 (1 × 50 bp SR, v3) platform.

### Small RNA-seq data processing and mapping

Obtained demultiplexed raw sequencing reads (fastq) were processed by cutadapt v1.9 (16) which was used to trim adapter sequences and low quality bases (>Q20), and to discard sequences shorter than 18 nucleotides in length, with the following parameters for TruSeq data: “cutadapt -a TG-GAATTCTCGGGTGCCAAGG -m 18 -q 20 –discard-untrimmed”; and with the following parameters for NEXTflex data: “cutadapt -u 4 -a NNNNTGGAATTCTCGGGTGCCAAGG -m 18 -q 20 –discard-untrimmed”. The processed reads were then mapped to miRNA sequences from miRBase v22 (17) using mirAligner (18) with default parameters (1 mismatch, 3 nt in the 3’ or 5’ trimming variants and the 3 nt in 3’ -addition variants). The R package isomiRs v1.10.1 (19) with default parameters was used to generate the count matrix of miRNA counts per library. Samples with fewer than one million mapped reads were excluded from further analysis. The sequencing depth of mapped reads to miRNA reference sequences is shown in **Supplementary Figure 1**. Raw sequencing reads and quantified read-count data have been deposited at NCBI Gene Expression Omnibus (GEO) (20) under the accession number GSE138318.

### Blocking efficiency estimation

In order to estimate the efficiency of blocking oligos for miR-486-5p and miR-451a, read counts were normalized to counts per million (CPM) by employing the cpm() function from the R package edgeR (21). The blocking efficiency for each target miRNA (i) in each paired sample (j) was then calculated using the following formula:

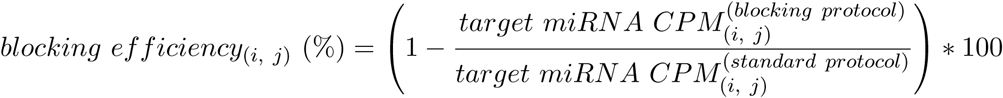

A one-sided Wilcoxon rank sum test was used to compare blocking efficiencies between NEXTflex and TruSeq protocols by employing wilcox.test() function from the base R package.

### Detectability determination

In order to compare miRNA detectability with or without the use of the blocking oligos, the obtained read counts were normalized to CPM as described previously and only the miRNAs which had at least 1 CPM in at least 75% of the libraries per protocol were considered as detected with confidence. The overlaps of detected miRNAs among different protocols were calculated and visualized using the R package ggupset (22). The simulation of random down-sampling of miRNA counts was performed by employing drarefy() function from the R package vegan (15), which returns probabilities for each miRNA to be detected in a random subsample. For each sample, miRNA counts were subsampled to seven different levels (5M, 4M, 3M, 2M, 1M, 0.5M and 0.1M). In the down-sampling experiment, a miRNA was considered to be detected when the probability was above 0.9.

### Blocking effect on quantitative performance estimation

The analysis of blocking oligo effect on non-targeted miRNA quantitative estimates was based on samples for which both blocked (modified) and unblocked (standard) libraries were generated. Pearson correlation analysis between blocked and unblocked paired samples was performed on the log-transformed (using pseudo-count of 1) CPM values of detected miRNAs. Pearson’s correlation coefficients were calculated using the cor() function (implemented in the R base package) with all observations except those of miR-486-5p and miR-451a. The R package DESeq2 (23) with default parameters and paired sample design was used to estimate the differential expression of miRNAs between blocked and unblocked small RNA libraries. The P-values resulting from Wald tests were corrected for multiple testing according to Bonferroni method (24). The miRNAs with a corrected P-value < 0.01 and |log2FC| >1 were considered to be significantly differentially expressed. The RNA-cofold algorithm from the Vienna RNA package (25) was employed to evaluate nonspecific hybridization between blocking oligonucleotides and down-regulated non-targeted miRNAs. All the data was visualized using the R package ggplot2 (26).

## Results

### Erythropoietic miR-486-5p and miR-451a domination reduces the detectability of low abundant miRNAs in whole blood samples

Human blood is a liquid, composite biological tissue consisting of multiple cells (erythrocytes, monocytes, neutrophils, lymphocytes, thrombocytes, etc.) and cell-derived components (platelets, EVs, etc.) suspended in a medium known as plasma (27). Besides blood-cell expressed miRNAs, blood also contains circulating miRNAs that are detected in serum, plasma or EVs (28, 29, 30). Therefore, the expectation is that the small RNA-sequenced whole blood sample should contain convolved information about the miRNA composition of all cellular and non-cellular blood components.

To test this hypothesis, we performed a compositional analysis of miRNA species in blood compounds and whole blood samples using blood miRNA catalog data (14). Here, for each sample, we calculated the Shannon diversity index, which combines the information about miRNA richness (number of detected different miRNA species) and evenness (proportion of each miRNA) within a given sample (31). Unexpectedly, whole blood samples showed the lowest average miRNA diversity (corrected P-value range: 9.94×10^−110^ – 1.42×10^−07^; **Figure 1**) as well as the lowest average level of richness (data not shown) values among separately sequenced blood compounds, meaning that more miRNAs were detected in every single blood compound than in whole blood itself.

To understand what is causing low miRNA diversity in the whole blood samples, we looked at relative miRNA abundances and observed high domination of miR-486-5p and miR-451a molecules in the samples, where they both comprise about ∼80% of total mapped reads (**Figure 1**). In addition to whole blood, we observed that these miRNAs are also dominant in erythrocytes, serum and partly in EVs. Interestingly, miR-486-5p and miR-451a were previously shown to be involved in erythroid development (6), which partly explains why these miRNAs are so abundant in erythrocytes. Since erythrocytes and reticulocytes together comprise more than 90% of blood cells, these miRNAs are even more abundant in whole blood. A high abundance of these two erythropoietic miRNAs in EVs and, especially, in serum might be explained by contamination due to hemolysis, which may occur during sample handling.

Taken together, these observations suggested that the domination of erythropoietic miR-486-5p and miR-451a in the whole blood small RNA libraries exhausts sequencing space and reduces the detectability of other low abundant miRNAs coming from less copious cells or non-cellular blood compounds.

### Erythropoietic miRNA blocking in 5’ ligation-dependent small RNA libraries

To deplete erythropoietic miR-486-5p and miR-451a, we designed linear oligonucleotides covering the longest stable complementarity of target miRNA sequence variants and bearing terminal modifications to prevent these oligonucleotides from participating in ligation reactions or being extended by polymerase. To design the oligonucleotides, we have obtained all sequence variants (isomiRs) of mature miR-486-5p and miR-451a molecules from our previously published whole blood smRNA-seq dataset (14). To identify the consensus sequences for each target miRNA, we calculated nucleotide frequencies at each position in sequence alignments of the precursor miRNAs and flattened them into one sequence (**Figure 2A**). The consensus sequences revealed higher variability within 3’ than within 5’ end of target miRNA sequences. This occurs due to non-templated nucleotide additions, which, beside cleavage-directed 5’ and 3’ end modifications, introduce additional variation within 3’ ends of miRNA sequences (32). Therefore, because of this miRNA feature, we decided to block 5’ end of target miRNAs and to prevent them from 5’ adapter ligation. For each target miRNA, we generated reverse complement oligonucleotides of the consensus sequence containing the most stable nucleotides starting from the 5’ ends of the molecules. Finally, on the 3’ ends of the oligonucleotides, we added C3 spacer (propyl group) modification, which prevents the blocking oligonucleotides from self-ligation and extension. Complementary DNA oligos bearing this modification have been previously shown to prevent 5’ adapter ligation to Drosophila 2S rRNA in smRNA-seq libraries (11).

In order to test our blocking oligos targeting miR-486-5p and miR-451a, we chose two 5’ adapter ligation-dependent methods, the commercially-available TruSeq and NEXTflex protocols. TruSeq is one of the most commonly used methods for small RNA library preparation employing adapters with invariant sequences, while NEXTflex utilizes adapters containing four degenerate nucleotides at the ligation ends as a strategy to reduce ligation bias, which is a well-known issue in small RNA library preparation (33).

The standard TruSeq protocol can be summarized in three core steps: 3’ adapter ligation, 5’ adapter ligation and reverse transcription (RT) accompanied by amplification of the cDNA library. Since the blocking oligos were designed to target 5’ ends of complementary miRNA sequences, in the modified TruSeq protocol, we have introduced these oligos prior to 5’ adapter ligation step (**Figure 2B**), where the complementary segments of targeted miRNA sequences and the blocking oligos take part in the formation of double-stranded RNA:DNA hybrids. These blunt-ended or slight 3’ DNA overhang-having double-stranded hybrids are not suitable substrates for T4 RNA ligase 1 to join a single-stranded adapter to the 5’ end of RNA strand in the hybrid. The hybrids without adapter sequences cannot be amplified and are therefore removed from the final library.

The standard NEXTflex protocol, besides having randomized adapter ends, also includes an additional step called 3’ adapter inactivation. In this step, end-filling is performed to fill the gaps of single-stranded 5’ overhang portions of pre-annealed 3’ adapters (34). As in the case of “blocked” RNA:DNA hybrids, the 5’ end-filled and blunt-ended adapter-oligonucleotide duplexes are poor substrates for T4 RNA ligase 1, which leads to reduced adapter dimer formation during the 5’ adapter ligation step. Due to the 3’ adapter inactivation step, we could not introduce our blocking oligos prior to 5’ ligation, because in order to anneal the blocking oligos to target miRNAs, the temperature has to be increased to at least 70 *°*C which could denature 3’ adapter-oligonucleotide duplexes with random nucleotide ends and lead to reduced library output. Therefore, in the modified NEXTflex protocol, we introduced and annealed our blocking oligos directly to total RNA samples prior to library preparation (**Figure 2C**).

### Blocking oligonucleotides with high efficiency reduce miR-486-5p and miR-451a sequences in small RNA libraries

In order to test the efficiency of our blocking oligos targeting the erythropoietic miRNAs, for each replicate sample, we generated unblocked (standard) and blocked (modified) libraries using both TruSeq and NEXTflex protocols. With this design, we generated 8 paired libraries for TruSeq and 5 paired libraries for NEXTflex using whole blood total RNA as an input. As expected, within the libraries that we generated by unmodified protocols, miR-486-5p was the most abundant miRNA in each library independent of the kit. On average, we obtained around ∼954K and ∼435K counts per million (CPM) of miR-486-5p using unblocked TruSeq and NEXTflex protocols, respectively. When using blocking oligos, we were able to suppress the average CPM values of miR-486-5p down to ∼21K and to ∼5K in TruSeq and NEXTflex protocols, respectively (**Figure 3A**). In contrast to miR-486-5p, we observed high variability of miR-451a expression within whole blood RNA samples which were prepared using unblocked NEXTflex protocol. The CPM values of this miRNA were also higher in NEXTflex than in TruSeq unblocked libraries. On average, we obtained ∼6K and ∼66K CPM of miR-451a using unblocked TruSeq and NEXTflex protocols, respectively. When using blocking oligos for miR-486-5p, the average CPM values were suppressed down to ∼150 and to ∼426 CPM in TruSeq and NEXTflex protocols, respectively (**Figure 3A**).

**Figure 3:**
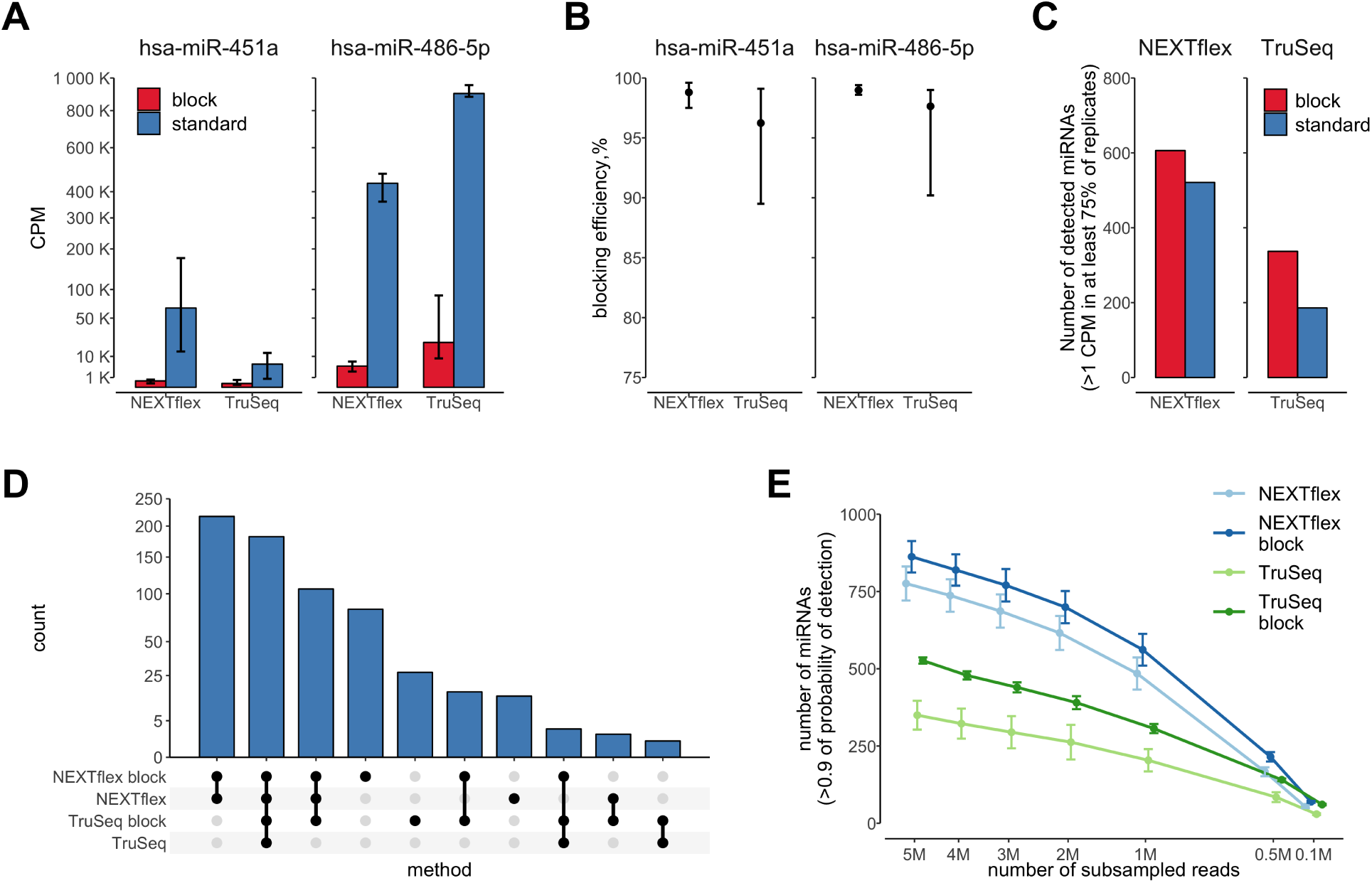
Blocking oligonucleotides efficiently suppress miR-486-5p and miR-451a, and increase detectability of other miRNA species in small RNA libraries. (**A**) A bar chart represents quantitative estimates of miR-486-5p and miR-451a in blocked (red) and unblocked (blue) libraries prepared by NEXTflex and TruSeq protocols. The y-axis depicts counts per million (CPM) and it is scaled by square root. The error bars indicate min and max values obtained from replicates. The graph illustrates a high degree suppression of miR-451a and miR-486-5p sequences in blocked NEXTflex and TruSeq libraries; (**B**) A dot plot represents blocking efficiencies (y-axis) of miR-486-5p and miR-451a in NEXTflex and TruSeq libraries. The data points depict mean values, whereas the error bars indicate min and max values obtained from replicates. The overall blocking efficiency is observed to be slightly better in the NEXTflex than in the TruSeq small RNA libraries (Wilcoxon P-value = 0.012); (**C**) A bar chart shows number of detected miRNA species (y-axis) in blocked (red) and unblocked (blue) libraries prepared by NEXTflex and TruSeq protocols. The detectability of miRNAs is increased in both blocked NEXTflex and TruSeq libraries; (**D**) An upset plot representing intersection of uniquely detected miRNA species amongst the set of the four protocols. The highest numbers of uniquely detected miRNAs are found in blocked libraries; (**E**) A line chart illustrates number of detected miRNAs (y-axis) in subsamples (x-axis; scaled by square root) of down-sampled libraries prepared by different protocols. The data points represent mean values, whereas the error bars depict standard errors of the mean. The graph displays a steady increase of detected miRNAs over the increasing size of subsampled miRNA counts.

By calculating blocking efficiencies, we observed that the blocking performance of linear oligonucleotides was slightly better in the NEXTflex than in the TruSeq protocol (P-value = 0.012). The blocking efficiency of the oligonucleotides for miR-486-5p ranged from 90.2% to 99.0% with an average of 97.7% in TruSeq, and from 98.6% to 99.4% with an average of 98.9% in NEXTflex protocol. In case of miR-451a, the blocking efficiency ranged from 89.5% to 99.1% with an average of 96.2% in TruSeq, and from 97.5% to 99.6% with an average of 98.8% in NEXTflex protocol (**Figure 3B**).

### Blocking oligonucleotides increase the detectability of low abundant miRNAs in blood-derived RNA samples

To test whether the depletion of erythropoietic miRNAs increases the information in blood-derived small RNA libraries, we compared miRNA detectability in blocked and unblocked libraries prepared by using TruSeq and NEXTflex methods. In the analysis, we considered a miRNA as detected if its CPM value was higher than 1 in at least 75% of the libraries prepared by exactly the same protocol. We detected the highest number of miRNAs in NEXTflex blocked libraries (n = 606), followed by NEXTflex unblocked (n = 521), TruSeq blocked (n = 337) and TruSeq unblocked (n = 186) libraries (**Figure 3C**). Interestingly, when looking at the overlapping and uniquely detected miRNAs, we not only observed uniquely detected miRNAs in the blocked protocols, but we were also able to detect several (n = 14) unique miRNAs in the unblocked NEXTflex protocol (**Figure 3D**). As expected, we found that most of the unique miRNAs were detected in the blocked NEXTflex (n = 82), followed by blocked TruSeq (n = 27) protocols.

To evaluate if the blocking effect on miRNA detectability is stable across libraries at varying sequencing depths, we have performed down-sampling of total mapped reads. We found that the usage of the blocking oligos already at 1 million subsampled reads increases miRNA detectability on average by 33.2% in the TruSeq and by 11.4% in the NEXTflex protocols. The increase of detectability stays more or less stable for both TruSeq (mean: 33.2%; range: 33–34%) and NEXTflex (mean: 11.4%; range: 10–14%) protocols, even when the subsample size is steadily increased to 5 millions of reads (**Figure 3E**).

Overall, these results clearly show that blocking oligos for miR-486-5p and miR-451a, independent of library size, increase detectability of other miRNA species in whole blood small RNA libraries.

### Blocking oligonucleotides have positive and some negative effects on non-targeted miRNA quantitative estimates

To evaluate the effect of miR-486-5p and miR-451a blocking on non-targeted miRNA species, we compared quantitative estimates of blocked and unblocked paired small RNA libraries which were generated using TruSeq and NEXTflex protocols. By looking at the distribution of average log-transformed CPM values of miRNAs, we observed that erythropoietic miRNA depletion resulted in a noticeable shift towards higher values in the density curves of libraries prepared by both TruSeq and NEXTflex protocols, which means that the CPM values of not only lowly but also of highly abundant miRNAs were increased proportionally. This global shift of log-transformed CPM values was more pronounced in the blocked TruSeq than in the blocked NEXTflex protocols (**Figure 4A**). We also observed consistent results when we looked at the paired blocked and unblocked libraries of each sample separately, where we saw an increase in the number of detectable miRNAs as well as a global increase of measured miRNA expression in the blocked libraries of both TruSeq and NEXTflex methods (**Figure 4B**). For both of the library preparation methods, on average, we observed a high concordance of miRNA expression (log-transformed CPM) estimates between paired blocked and unblocked libraries which was 0.94 (range: 0.80–0.97) for TruSeq, and 0.93 (range: 0.91–0.96) for NEXTflex protocols, in terms of Pearson correlation coefficient (**Supplementary Figure 2**).

**Figure 4:**
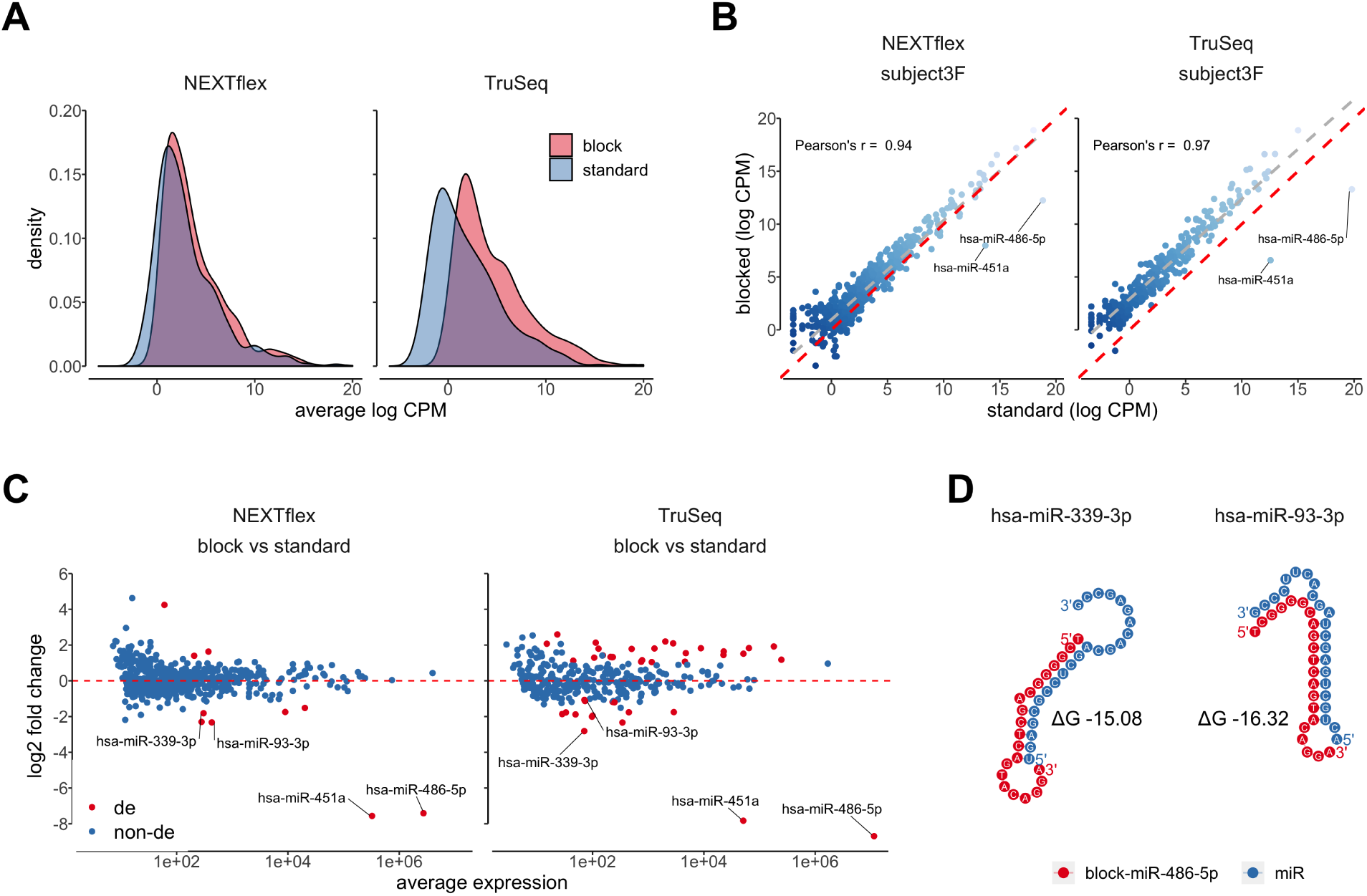
Blocking oligonucleotides for miR-486-5p and miR-451a have positive and some negative effects on measured expression of non-targeted miRNAs. (**A**) A density chart shows distributions of averaged log2-transformed CPM values of blocked (red) and unblocked (blue) libraries prepared by NEXTflex and TruSeq protocols. The expression values of each miRNA were averaged per protocol (x-axis). The graph illustrates a proportional global shift of log2-transformed CPM values towards higher expression estimates in both blocked NEXTflex and TruSeq libraries; (**B**) A scatter plot represents correlation of log2-transformed CPM values of paired blocked (y-axis) and unblocked (x-axis) exemplary libraries generated by NEXTflex and TruSeq protocols. The red dashed line divides panel in two equal parts, whereas the grey dashed line displays a linear regression curve. The chart illustrates a high concordance of miRNA expression values between blocked and unblocked paired libraries; (**C**) An MA plot shows paired-differential expression analysis results of blocked versus unblocked libraries, where log2 fold changes are presented on the y-axis and averaged normalized counts on the x-axis. The red colored dots indicate significantly differentially expressed miRNAs (corrected P-value < 0.01; absolute value of log2 fold change > 1). The miRNAs which were found to be down-regulated in both NEXTflex and TruSeq libraries are labeled with miRNA names; (**D**) A dot chart represents predicted non-specific interactions between miR-486-5p blocking oligonucleotide (red) and two non-targeted miRNA molecules (blue). These interactions might cause decreased expression of miR-339-3p and miR-93-3p in blocked small RNA libraries.

To obtain further insights whether the blocking oligos may have an impact on measured expression of non-targeted miRNAs, for TruSeq and NEXTflex methods separately, we performed differential expression analysis between blocked and unblocked libraries using DESeq2 with paired sample design. As expected, we saw a global increase of log-transformed fold change values, which was generally higher for low abundant miRNA species in both comparisons of blocked versus un-blocked paired libraries (**Figure 4C**; **Supplementary Tables 1 and 2**). We also identified significantly up-regulated high abundant miRNAs, especially in the libraries prepared by TruSeq protocol, which might be explained by 5’ ligation bias, because, in the absence of the highly abundant targeted-miRNAs, non-target miRNAs might have different ligation efficiency resulting in a non-proportional increase.

In addition to miR-486-5p and miR-451a, unexpectedly, we also observed 14 significantly down-regulated (corrected P-value < 0.01; absolute value of log2 fold change > 1) miRNAs in TruSeq, and 5 miRNAs in NEXTflex libraries. To see whether there is a systemic problem with blocking oligos and down-regulated miRNAs, we looked at the miRNAs which were commonly down-regulated in both methods, and besides miR-486-5p and miR-451a, found two such molecules: miR-339-3p and miR-93-3p (**Figure 4C**). By using RNA cofold analysis we showed that these two miRNAs may interact with blocking oligos of miR-486-5p and form secondary structures which might inhibit or reduce 5’ ligation of the non-targeted miRNAs (**Figure 4D**).

Overall, in addition to a number of positive effects such as increased miRNA detectability, global increase of expression values and little effect on non-targeted miRNA species, negative effects of the blocking oligos such as non-specific hybridization may also appear. This should be taken into consideration when analyzing the data.

## Discussion

The erythroid-specific, highly dominant and less likely informative miR-486-5p and miR-451a transcripts are muting detection of lowly abundant miRNAs in whole blood-derived small RNA libraries. To overcome this problem, we have developed a cost-effective and easy-to-use hybridization-based protocol to deplete these erythropoietic miRNAs from small RNA libraries. This method, besides custom oligos, does not require any other additional reagents, and is easily compatible and adjustable with 5’ ligation-based small RNA library preparation methods such as Illumina’s TruSeq and Perkin Elmer’s NEXTflex protocols.

We demonstrate the high overall blocking efficiency of our oligonucleotides, whereas, the average efficiency was slightly better in NEXTflex (98,9%) than in TruSeq (96,9%) libraries. As a consequence of erythropoietic miRNA blocking, the measured expression as well as detectability of low abundant miRNA species was considerably increased. This increase was more pronounced in the TruSeq than in the NEXTflex libraries, probably due to higher domination of miR-486-5p in the unblocked TruSeq libraries. It seems that this particular miRNA is highly preferred by TruSeq method (∼90% of total mapped reads) and once it is depleted, much more space is freed up for other miRNA species. Even though the relative increase of detectability is higher in blocked TruSeq libraries, the nominal detectability is much higher in the blood RNA-derived NEXTflex libraries, which bear more miRNA species than TruSeq libraries even without the use of blocking oligos. This is probably due to utilization of degenerate adapters which reduce ligation bias in small RNA libraries.

We also demonstrate that our oligonucleotides do not reduce the reproducibility of the quantitative estimates of non-targeted miRNAs and, moreover, does not remove or significantly disturb individual-specific biological variation. In terms of Pearson correlation coefficient, the reproducibility of blocked and unblocked libraries are very similar in both TruSeq and NEXTflex libraries. Despite good performance of our hybridization-based method, we also detected some off-target effects, which were observed independent of library preparation method, suggesting a systemic effect of blocking oligos on at least two non-targeted miRNAs. On the other hand, since it is a systemic effect, in studies such as differential expression between cases and controls this effect should even out.

In the current version of the protocol, we have designed and optimized the blocking oligonucleotides and their hybridization conditions specifically for miR-486-5p and miR-451a; however, the oligo design principles can be adapted to target any other miRNA molecule. Of note, each tissue or cell type might contain different isomiR composition (35), and therefore, we would recommend to ensure that the designed oligo covers all nucleotides at the 5’ end of the target sequence. This can be achieved by mapping reads to miRNA isoforms or alternatively, by simply extending the 5’ end of canonical miRNA sequence a few nucleotides upstream of the reference position. Even though this recommendation is not tested in the current design of the study, it is well-known that T4 RNA ligase 1 prefers single stranded RNA as a substrate (36, 37) and therefore, 5’ overhang of acceptor RNA in “blocked” RNA:DNA hybrid might result in adapter ligation and in this way may reduce blocking efficiency of the oligonucleotides.

We demonstrated compatibility of this method with TruSeq and NEXTflex protocols; however, theoretically the blocking oligos should be also compatible with other 5’ ligation-based methods which employ T4 RNA ligase 1 (such as NEBNext, QIAseq, CleanTag, etc.) to attach an adapter oligonucleotide to the 5’ end of small RNA molecules.

## Supporting information

Supplementary Protocol

## Notes

https://github.com/ikmb/erythromiR_depletion_analysis

