## Supplementary Protocol for "Depletion of erythropoietic miR-486-5p and miR-451a improves detectability of rare microRNAs in peripheral blood-derived small RNA sequencing libraries"

### SUPPLEMENTARY PROTOCOLS

### 1. Preparation of blocking oligonucleotide mix

Required materials:

1. Custom DNA oligo 1 (IDT);
2. Custom DNA oligo 2 (IDT);
3. TE buffer (Ambion).

**Table 1:** Features of the blocking oligonucleotides. The oligonucleotides were designed based on sequences from miRBase v22 database.

| Name | Oligonucleotide sequence (5' to 3') | Targeted-microRNA | Tm $^o$C (DNA/RNA) |
|---|---|---|---|
| Oligo 1 | TCGGGGCAGCTCAGTACAGGA/3SpC3/ | hsa-miR-486-5p | 62.5 +/- 2.7 |
| Oligo 2 | ACTCAGTAATGGTAACGGTTT/3SpC3/ | hsa-miR-451a | 51.5 +/- 2.7 |

Preparation:

1.1. Resuspend each oligonucleotide in TE buffer to achieve a 100 $\mu$M concentration.
1.2. Combine Oligo 1 and 2 solutions at even volumes to prepare stock solution of the blocking oligo mix.

### 2. TruSeq Small RNA Library Prep (#15004197 v02) protocol modification

The only modified step of the protocol is "Ligate 3' Adapter", where the blocking oligos are introduced to reaction mix of each sample. The rest of the steps follow the standard protocol.

Additional preparation before starting the library preparation:

1. Dilute blocking oligo mix (from step 1.2) in TE buffer to 20 $\mu$M solution.

#### *Alternative Ligate 3' Adapter (page 12)*

14. ...
15. Remove from the thermal cycle and place on the ice block. Add 1 $\mu$l of blocking oligo mix (20 $\mu$M) directly to the reaction.
16. Place on the thermal cycle and incubate at the following:

- 90 $^o$C for 30s;
- Ramp from 65 to 45 $^o$C at a speed of 0.1$^o$C per sec.
- 4 $^o$C for at least 30s.

17. Immediately proceed to "Ligate 5' adapter" step of the standard protocol.

### 3. NEXTFLEX Small RNA-Seq Kit v3 (V19.01) protocol modification

The only modified step of the protocol is "Step A", where the blocking oligos are introduced to total RNA of each sample. The rest of the steps follow the standard protocol.

Additional preparation before starting the library preparation:

1. Dilute blocking oligo mix (from step 1.2) in TE buffer to 10 $\mu$M solution.

#### *Alternative Step A (Page 9)*

1. For each sample, combine the following reagents on ice in nuclease-free 96-well PCR plate:

_ $\mu$l RNA
_ $\mu$l Nuclease-free Water
1 $\mu$l blocking oligo mix (10 $\mu$M)

---

10.5 $\mu$l TOTAL

2. Place on the thermal cycle and incubate at the following:

- 90 $^o$C for 30s;
- Ramp from 65 to 45 $^o$C at a speed of 0.1$^o$C per sec.
- 4 $^o$C for at least 30s.

3. Place on ice and proceed to the next step of the standard protocol.
4. ...
