## Supplementary Figures for "Depletion of erythropoietic miR-486-5p and miR-451a improves detectability of rare microRNAs in peripheral blood-derived small RNA sequencing libraries"

---

### SUPPLEMENTARY FIGURES

---

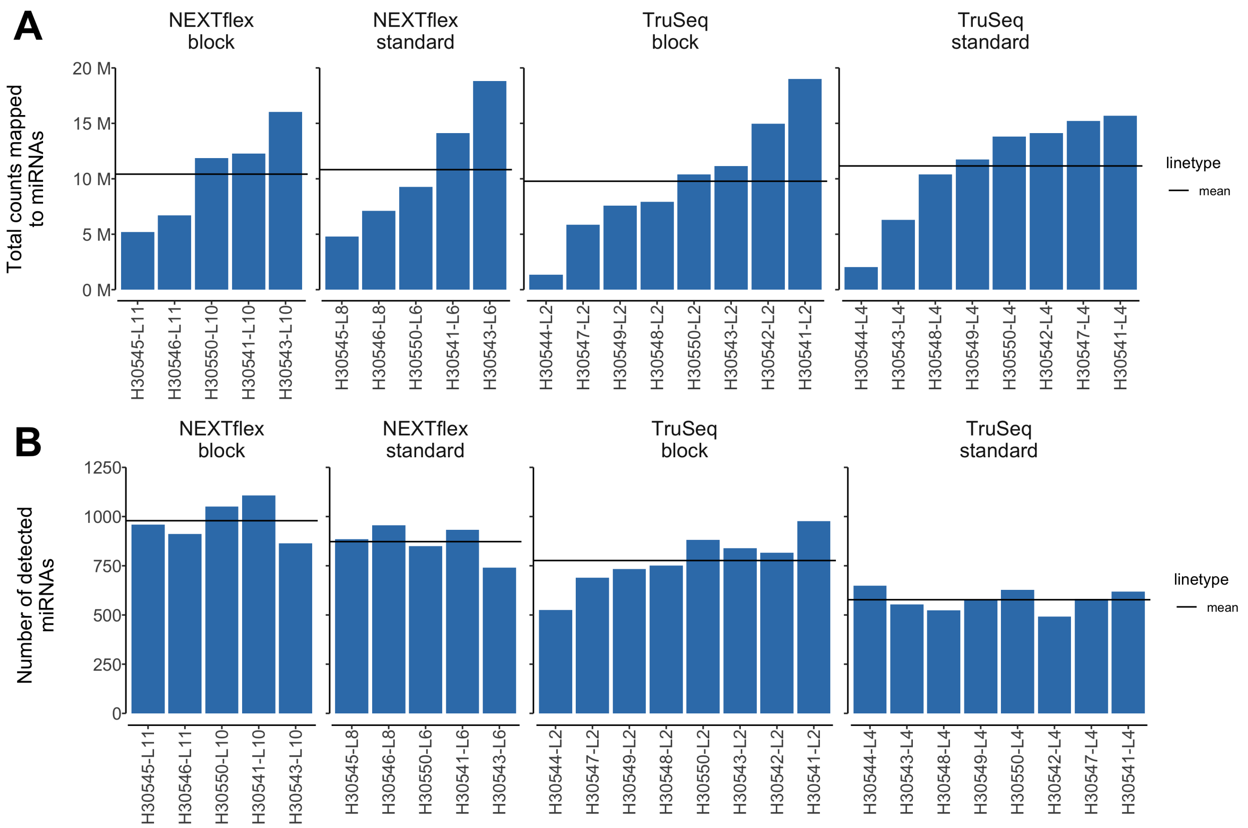


**Supplementary Figure 1: A sequencing depth of mapped reads with reference to miRBase version 22 database.** (**A**) A bar chart showing number of mapped reads (in million) to miRNA hairpins (y-axis) of each small RNA library (x-axis); (**B**) A Bar chart representing number of uniquely detected (at least 1 read) miRNAs (y-axis) in every small RNA library (x-axis). Solid line represents average value per set of the four protocols. While the average library size (total reads mapped to miRNAs) values of all the libraries are similar, the average numbers of uniquely identified miRNAs are higher in the blocked libraries.

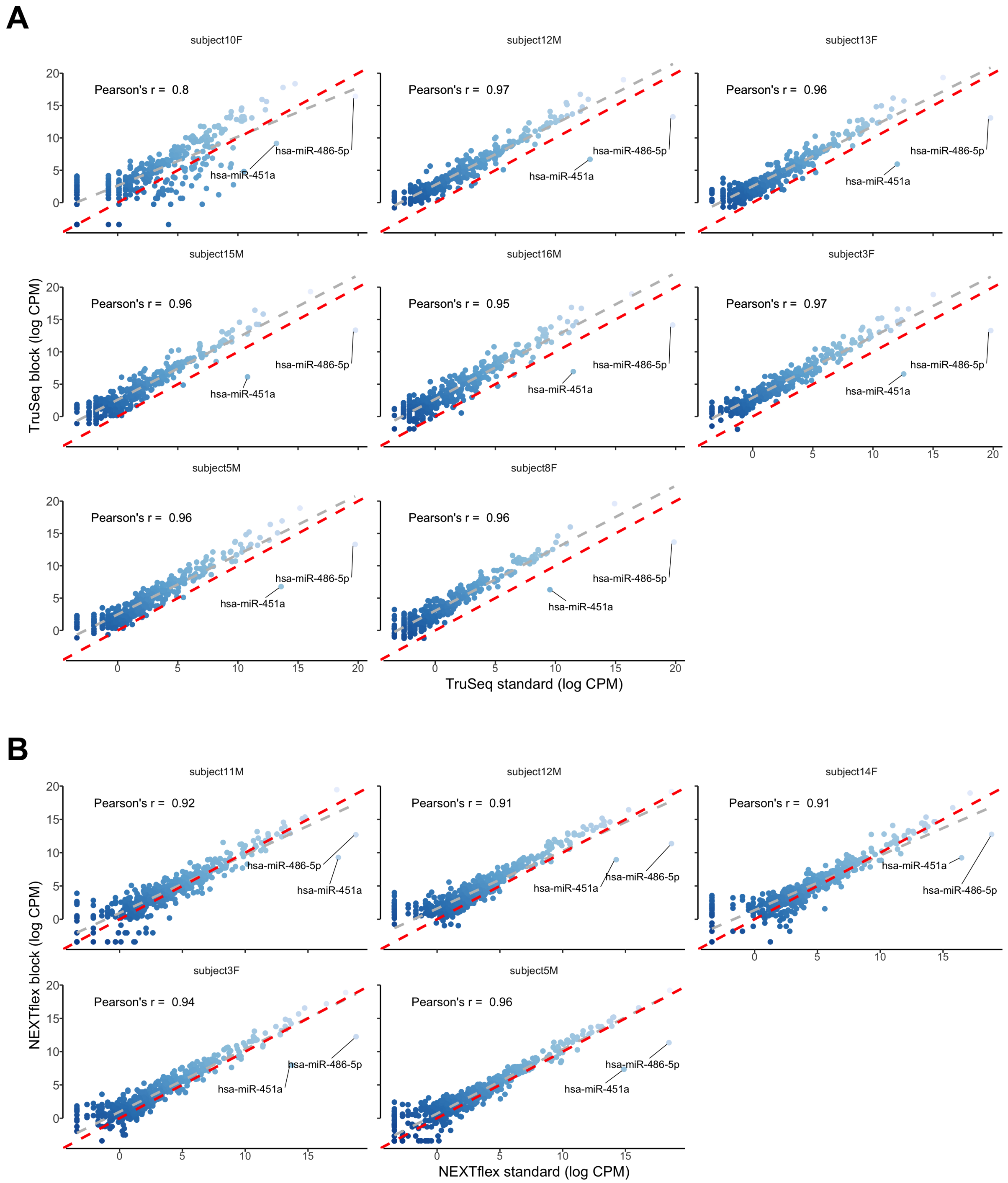


**Supplementary Figure 2: A concordance of miRNA expression estimates between paired blocked and unblocked libraries.** A scatter plot represents correlation of log2-transformed CPM values of paired blocked (y- axis) and unblocked (x-axis) exemplary libraries generated by TruSeq (**A**) and NEXTflex (**B**) protocols. The red dashed line divides panel in two equal parts, whereas the grey dashed line displays a linear regression curve. The chart illustrates a high concordance of miRNA expression values between blocked and unblocked paired libraries.
